# Stopping Speed in Response to Auditory and Visual Stop Signals Depends on Go Signal Modality

**DOI:** 10.1101/2024.01.10.575126

**Authors:** Simon Weber, Sauro E. Salomoni, Rebecca J. St George, Mark R. Hinder

## Abstract

Past research has found that the speed of the action cancellation process is influenced by the sensory modality/modalities of the environmental change that triggers it. However, the effect on selective stopping processes (where participants must cancel only one component of a multi-component movement) remains unknown, despite these complex movements often being required as we navigate our busy modern world.

Thirty healthy adults (mean age = 31.1 years, SD = 10.5) completed five response-selective stop signal tasks featuring different combinations of “go signal” modality (the environmental change baring an imperative to initiate movement; auditory or visual) and “stop signal” modality (the environmental change indicating that action cancellation is required; auditory, visual, or audiovisual). Electromyographical (EMG) recordings of effector muscles allowed detailed comparison of the characteristics of voluntary action and cancellation between tasks.

Behavioural and physiological measures of stopping speed demonstrated that the modality of the *go* signal influenced how quickly participants cancelled movement in response to the stop signal: stopping was faster in two cross-modal experimental conditions (auditory go - visual stop; visual go - auditory stop), than in two conditions using the same modality for both signals. A separate condition testing for multisensory facilitation revealed that stopping was fastest when the stop signal consisted of a *combined* audiovisual stimulus, compared to all other go-stop stimulus combinations. These findings provide novel evidence regarding the role of attentional networks in action cancellation and suggest modality specific cognitive resources influence the latency of the stopping process.

## Action Cancellation in Response to Visual and Auditory Stimuli

Response inhibition, our ability to cancel movements in response to changes in the environment, represents a key aspect of executive function. This ability is commonly assessed in the laboratory using a behavioural paradigm known as the stop signal task. Stop signal tasks are mostly comprised of “go trials”, which require the participant to enact a prescribed movement (e.g., a button press) in response to a “go signal”. Randomly interspersed amongst go trials are a small proportion of “stop trials”. Stop trials also feature a go signal, though this is followed by a “stop signal” which indicates that the participant must cancel their prepotent response. This paradigm has been broadly used for clinical and non-clinical research in psychology, biology, psychiatry and neuroscience (Verbruggen et al., 2019).

Notably, past research has found stopping speed is influenced by the modality of the stop signal; with a number of papers noting that stopping is *faster* in response to auditory stimuli than visual stimuli (Morein-Zamir & Meiran, 2003; Ramautar et al., 2006; Van Der Schoot et al., 2005). Research using electroencephalography (EEG) has suggested that this observation is primarily a result of bottom up differences (i.e., faster auditory *c*.*f*. visual processing) rather than differences in top-down inhibitory processes (Carrillo-de-la-Peña et al., 2019).

However, faster stopping to auditory cues is not a consistent finding (Ikarashi et al., 2022). This is likely due to heterogeneity in behavioural tasks. For instance, research by Ikarashi et al. (2022), used an auditory go signal whereby the frequency of the tone (1000Hz or 1500Hz) indicated whether a le- or right-hand response should be made. On stop trials, the same frequency tone was then played, following a delay, to indicate that this selected response should be inhibited. No significant differences in stopping speed were noted when this audio condition was compared to either a condition using visual go and stop signals (white and red arrows) or haptic stimuli (electrical pulses delivered to the selected/to-be-stopped index fingers; 200μs at 2.5x the participant’s sensory threshold, for both go and stop signal). Notably, the aforementioned papers that report faster stopping speed in response to auditory, compared to visual, stop signals all used visual go signals (Carrillo-de-la-Peña et al., 2019; Ramautar et al., 2006; Van Der Schoot et al., 2005). Accordingly, the experiments tested differences in stopping speed between auditory and visual *stop* signals, but also (inadvertently) induced differences between conditions in regards to whether the go and the stop stimuli were presented in the same, or differing modalities (e.g. both visual, or visual go – audio stop). The current experimental conditions were specifically designed to tease apart the sources of faster stopping performance, investigate this difference by comparing multiple modality combinations of go and stop cues (i.e., visual or auditory).

### Selective Stopping and Stopping Delays

Often, reacting appropriately to new sensory information requires complex inhibitory responses, whereby only a subset of our current motor plans must be cancelled, while others continue. For example, if someone is climbing a ladder and notices a spider on the next rung (where they are about to place their hand) they must inhibit the movement of their arm, but continue with their current step, to secure their balance. The term “response selective inhibition” refers to movements that require selective cancellation of a subset of the effectors currently engaged in motor plans. Past research indicates that this type of action cancellation is performed “sub-optimally” as the ongoing movements which coincide with the action cancellation are typically delayed. In experimental settings, this phenomenon (the delay in the non-cancelled component of the movement) has historically been referred to as the stopping interference effect (for a review see Wadsley et al., 2022). Here we refer to this phenomenon using the term “*stopping delay*” to reflect recent research suggesting that, rather than a global stop *interfering* with a continued movement, the non-cancelled component instead represents a unique, re-programmed movement (Salomoni et al., 2023; Gronau et al., 2023).

The mediators of the stopping delay are a topic of current investigation. Both inhibition associated with attentional capture (Weber et al., 2023; Wessel & Aron, 2013) and mechanical delays involved in action re-programming are thought to be involved. Stopping delays may also be mediated in part by a transient global ‘pause’ which is distinct from the conscious cancellation of a movement (Diesburg & Wessel, 2021).

Selective stopping delays have only been investigated in response to stop signals presented in the visual medium. That is, no previous research has investigated response-selective action cancellation in response to *auditory* stop cues. As such, the degree to which stopping delays arise from processing in the prefrontal inhibitory networks mediating action cancellation (Aron, 2011), and the degree to which it is influenced by sensory-specific processing, is unknown. Given the many real word contexts which require selective inhibition in response to *auditory* stimuli, this is surprising. For example, if you are driving and an accompanying passenger notices a pedestrian crossing the road in your blind spot, their vocalisation indicates that inhibition is required; you must inhibit the movement of your foot, which was moving to press the accelerator, while continuing to turn the wheel, to prevent the car from hitting the curb. Or similarly, many modern cars include sensors which beep at the driver if they are about to reverse into an object. In these situations, the efficient coordination of limbs is vital, raising questions regarding how effectively humans can selectively inhibit movements in response to auditory environmental cues. Here, we seek to investigate how efficiently multicomponent movements can be inhibited in response to visual and auditory cues, assessing the degree to which stopping delays generalise across sensory modalities.

### Multisensory Facilitation and Response Inhibition

The phenomena of multisensory facilitation is manifested as faster reaction mesto a combination of simultaneous visual, auditory or haptic stimuli than to stimuli presented in a single sensory medium (Hershenson, 1962; Molholm et al., 2002). Interestingly, multisensory facilitation has not been observed with respect to the speed of the inhibitory process, i.e., action cancellation c.f. action execution (Stock et al., 2017). Furthermore, Strelnikov et al., (2021) observed that a combined audio-visual stop signal actually had a *deleterious* effect on stopping speed, relative to stopping speeds in response to a visual stimuli alone. To purportedly increase ecological validity, Strelnikov and colleagues used human faces as visual stimuli and human voices as auditory stimuli. To test whether their finding generalises to more generic/conventional stimuli, the current experiment included a condition with a combined auditory and visual stop signal featuring a colour change in the visual stimuli presented simultaneously with computer-generated tones.

### Using EMG to Determine Stopping Latency

Stopping speed is most commonly indexed by estimating stop signal reaction time (SSRT); this yields a single estimate of stopping speed per participant, per condition, based on behavioural data (Verbruggen et al., 2019). An alternative approach involves using a single-trial electromyographic (EMG) measure to determine stopping latency, known as CancelTime (Jana et al., 2020; Raud et al., 2022). Given that the validity of SSRT measures has been called into question as a result of violations to the model underlying them (Gulberti et al., 2014; Verbruggen & Logan, 2015), especially in selective stopping contexts (Bissett et al., 2021), we used both measures in the current paper. Using Cancel Time as a more precise measure of stopping speed, in conjunction with the conventional behavioural measure (SSRT), allowed for a more detailed comparison of stopping speeds in different sensory mediums than has been possible in previous research.

In sum, the current research used behavioural manipulations and electrophysiological measures to explore differences in action initiation and cancellation in selective stopping tasks using auditory and visual stimuli, with the following goals: **1)** Apply, for the first time, EMG-based measures of stopping speed (Canceltime) to determine whether sensory modality (i.e. auditory, visual) of the go and stop signals affects stopping speed; **2)** Compare selective stopping delays across different stimulus modalities to further understand the psychological and physiological basis of this delay; **3)** Assess whether multisensory facilitation of the inhibitory response is observed when an audiovisual stop stimulus is utilised c.f. an audio or visual stimulus alone.

## Methods

### Participants

A group of 31 healthy adults took part in the study, though the data from 1 participant was excluded due to very low stop trial success rates. Following exclusions, the data from 30 participants (mean age = 31.1 years, SD = 10.5, range = 18-53; 19 female, 11 male) was included. The sample size was determined based on our previous research which used cohorts of 24 and 23 and was sufficiently powered to detect small differences in CancelTime within these groups (Weber et al, 2023). We recruited sufficient participants to allow for the detection of similar effect sizes. Participants were recruited via the University of Tasmania School of Psychological Science’s research participation system. Participants received either two hours of course credit (if needed) or a $20 shopping voucher as compensation for their time. All participants reported that they had normal or corrected-to-normal vision and normal hearing in both ears. All participants provided informed consent prior to taking part in the experiment. This research was approved by the Tasmanian Human Research Ethics Committee.

### Procedure

#### Computer Interfaces

Participants were seated approximately 80cm from a 27 inch computer monitor with forearms and hands resting shoulder width apart on a desk. Hands were pronated, with the lateral aspect of each index finger rested against one of two custom made response buttons. Buton presses were registered via The Black Box Toolkit USB response pad and recorded using PsychoPy3 (Peirce et al., 2019). The buttons were mounted in the vertical plane, such that registering a response required participants to abduct their index fingers (towards the centre of the desk). As such, the first dorsal interossei muscle acted as the primary agonist; a muscle that is easily accessible for measuring contractions with electromyography (EMG). During the experiment, participants were required to press the button/s in response to auditory and visual stimuli. Auditory stimuli were generated using the free software Audacity and played through binaural headphones. The volume of the auditory stimuli was set at a comfortable but clear volume determined during piloting (∼40dB), which was below the volume of the startle reflex (Ludewig et al., 2003). Each participant was asked if they were comfortable with the volume (which was the same for all participants) and given the option to have it adjusted prior to commencing the experiment. No participants requested the volume be changed from the initial level. The total duration of the experiment was approximately 1.5 hours.

#### Electromyography (EMG)

EMG was recorded using disposable adhesive electrodes placed on the first dorsal interossei of participants’ le and right index fingers. On each hand, two electrodes were placed in a belly-tendon montage, with a third ground electrode placed on the head of the ulna. EMG data were recorded using the computer software Signal (Cambridge Electronic Design Ltd.). The analogue EMG signals were sampled at 2000Hz, amplified 1000 times and band-pass filtered between 20-1000 Hz. Participants were requested to keep their hands as relaxed as possible between trials, and only activate muscles to press the buttons. Once the experiment commenced, the researcher would remind the participant to relax their hands between trials, if real-time recordings (visual to the experimenter but not the participant) revealed background muscle activity.

### Behavioural Tasks

#### Overview

The experiment consisted of five conditions which were developed and delivered using Psychopy3 (Peirce et al., 2019), and which are overviewed schematically in Figure 1. Four of these conditions constitute all possible combinations of auditory and visual go and stop cues, specifically: **1**) Go-Visual-Stop-Visual **2**) Go-Visual-Stop-Audio **3**) Go-Audio-Stop-Visual and **4)** Go-Audio-Stop-Audio. To test for possible intersensory facilitation effects of stopping processes arising from a multisensory stop cue, one additional condition was included: **5**) Go-Visual-Stop-Audiovisual. With regards to task order, five conditions resulted in too many permutations of to complete a full counterbalancing sequence (5! = 120); accordingly, a Latin square design was utilised to minimise practice and fatigue effects. Each condition included 150 trials (comprised of 100 “go trials” and 50 “stop trials”; 33% stopping likelihood; described below). All trials, irrespective of condition or trial type, commenced with a fixation cross appearing for a variable duration drawn from a truncated exponential distribution ranging from 250-1000ms. This served to reduce the temporal predictability of the appearance of the subsequent go stimulus, thus ensuring participants were responding to the stimulus, rather than predicting its appearance (Verbruggen et al., 2019) and time-locking their response to the appearance of the stimulus.

**Figure 1:**
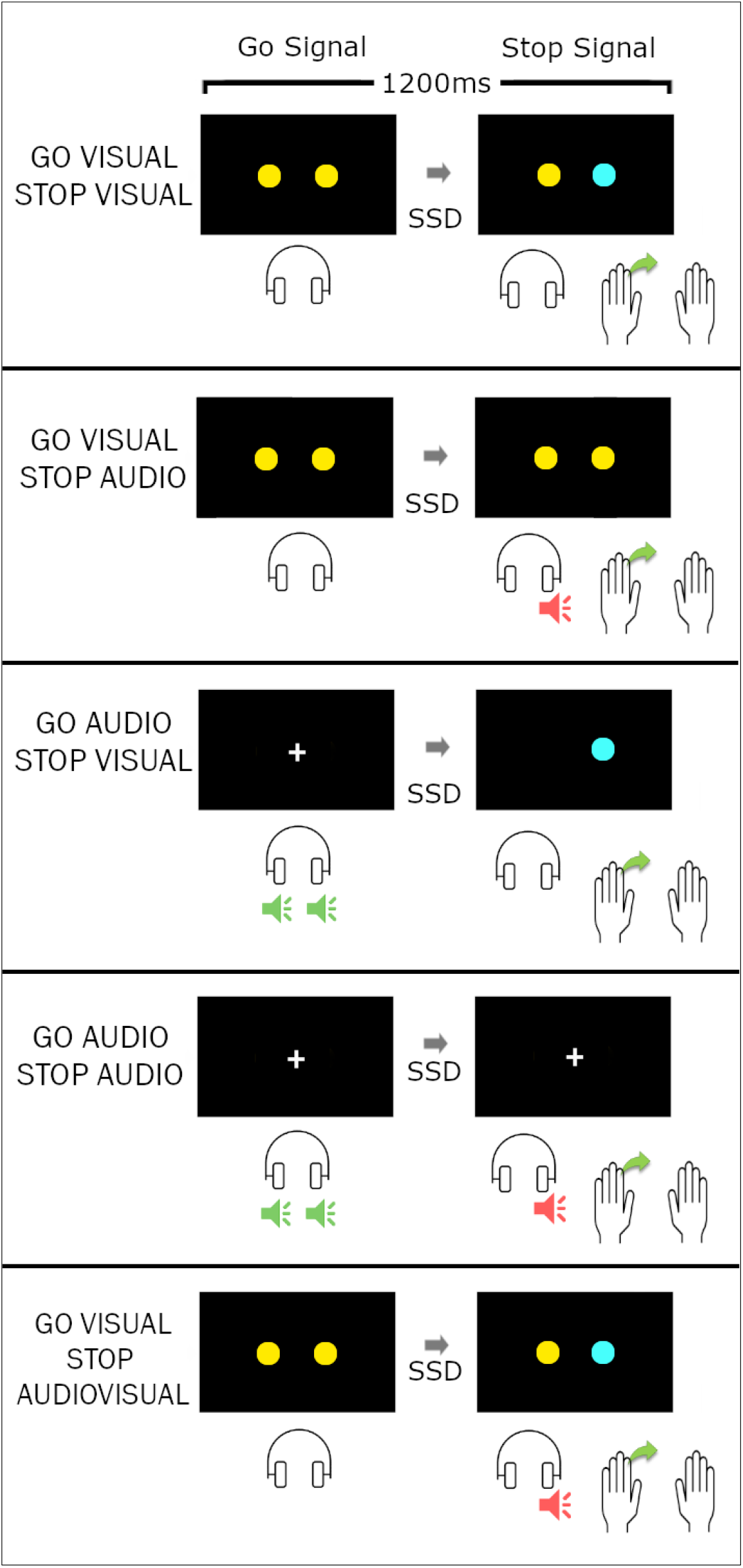
Trial sequence and timing of stop-right trials from each of the five main experimental conditions. Stop trials are shown as this allows for visual demonstration of the differences in go signal (the first pane in each row) and the stop signal (the second pane in each row) in each condition. Auditory go signals (pictured in green) involved a 220Hz sine wave played in both ears for 200ms, while auditory stop signals (pictured in red) involved a 440Hz sine wave played in one ear for 200ms. The hands are included to indicate the correct response (a button press with the left hand and no button press with the right hand).

#### Go Only Blocks and Go trials

Prior to completing the main experimental conditions, participants completed two short “go-only” blocks, each consisting of 15 go trials which required participants to enact a bimanual movement (press both buttons) in response to a go signal, which could be either visual or auditory depending on the condition. One go-only block involved go trials with auditory go signals and the other involved visual go signals. Auditory go signals used a 200ms duration tone (220Hz sine wave) played simultaneously through le and right headphone speakers. While this played, the fixation cross remained on the screen (so that the disappearance of the fixation cross couldn’t be used as a simultaneous visual indicator that the trial had begun). Visual go signals consisted of two yellow circles 3cm in diameter and 15 cm apart, appearing either side of the centre of the screen. From the onset of the go signal, the response window lasted 1200ms. This was followed by a feedback screen displaying the reaction time (in milliseconds) of both the le and right hands or, if participants failed to press both buttons, text reading “incorrect response”. The feedback screen lasted 1000ms before the screen was blank for 100ms and the next trial began.

Go trials during the main experimental conditions were similar to the go trials in the go-only blocks, with the one exception that additional feedback was used to minimise the strategic slowing of responses in anticipation of stop signals: The presence of stop signals can encourage proactive slowing, i.e., participants delay responding and wait for the stop cue, in an attempt to increase their chance of successfully stopping. To mitigate this strategy (which can invalidate measures used to assess stopping speed), if participants’ responses were more than 150ms slower than their average reaction times from the go-only blocks (where stop trials were not anticipated), additional feedback appeared above the RTs shown during the feedback screen saying, “You’ve slowed down”. The average RT calculated from the go-only blocks excluded very fast responses (<100ms) and asynchronous responses (difference of >50ms), to prevent anticipatory or accidental responses from influencing the threshold RT used for the feedback.

#### Stop Trials

One third of trials in each condition (50 per block) were stop trials, divided into “le-stop” trials (25 trials) and “right-stop” trials (25 trials). During stop trials the go signal appeared in the same manner as it did in go trials, though after a variable “stop signal delay” (SSD) it was followed by a stop signal. In conditions with visual stop signals (Figure 1) the stop signal was a blue circle appearing on either the le or right side of the screen. In conditions with auditory stop signals a tone (a 200ms duration 440Hz sine wave, easily distinguishable from the go signal) was played through either the le or right headphone speaker.^1^ Upon presentation of the stop signal, participants were required to inhibit their response on the side where the stop signal occurred, while continuing their response on the other side (that is, response-selective stopping was required). If participants succeeded in selectively cancelling their movement (e.g., pressed only the right button when the stop signal occurred on the le), text appeared during the feedback window reading “correct”. If participants made any other response (pressed both buttons, failed to press either button, pressed the button on the wrong side) the stop trial was counted as unsuccessful, and the feedback text displayed “incorrect”.

#### Stop Signal Delay (SSD)

A staircasing algorithm adjusted the timing of the SSD after each stop trial, separately for stop-le and stop-right trials. initially, both SSDs were set to 150ms, and this was adjusted based on success in the previous trial: following a correct stop-right trial the SSD for the subsequent stop-right would increase by 50ms, making it harder to stop on the next stop trial. Following a failed stop-right trial the SSD would decrease by 50ms, making it easier to stop on the next trial. Thus, the overall probability of successfully stopping was approximately 50% considering all stop trials in the condition (Verbruggen et al., 2019). The minimum and maximum SSDs were 50ms and 400ms, respectively.

#### Breaks and Trial Order

Aer every 50 trials, a short rest break was triggered. During the breaks, text appeared asking participants to rest their hands for a moment and to press one of the buttons when they were ready to continue. Below this text message, feedback was displayed indicating the proportion of correct stop trial responses and average reaction time of correct go trial responses from the previous 50 trials. For each condition, trial order was pre-randomised with the constraints that no more than two stop trials occurred consecutively and at least two go trials followed each break before a stop trial occurred. All conditions started with a series of instruction slides appearing on the screen explaining the task. Participants worked through these at their own pace, progressing through the slides with button presses, with the experimenter able to provide any additional clarification, if required.

#### Behavioural Data Analysis

Statistical analyses were conducted using generalised linear mixed models (GLMMs) and linear mixed models (LMMs). We ran these in the statistical packages of JASP (JASP Team, 2023) and Jamovi (The Jamovi Project, 2021), which operates via the statistical language R (R Core Team, 2021). GLMMs were initially run with a maximal random effects structure. If this failed to converge, a stepwise approach to model simplification was employed, first removing interaction terms from the random effects structure, followed by slope terms (Barr et al., 2013).

#### SSRT

SSRT was calculated using the integration method with replacement of omissions (Verbruggen et al., 2019). The model that underlies this calculation requires that mean RT on unsuccessful stops is faster than mean RT on go trials (Verbruggen et al., 2019). In the Go-Audio-Stop-Audio condition, data from two participants was excluded for not meeting this requirement, as well as data from one participant in the Go-Visual-Stop-Audio condition, and another from the Go-Visual-Stop-Visual condition. Given each participant only has a single SSRT score per condition, a general linear model (i.e., a repeated measures ANOVA) was run on this data with a single variable for condition, with five levels (please refer to Figure 1). Planned contrasts were used to assess the effects of stop-type and go-type. Another set of contrasts was used to check whether there was facilitation of the stop process in audiovisual stop signal condition compared to the Go-Visual-Stop-Visual and Go-Visual-Stop-Audio conditions (given that these two conditions are most reflective of the typical implementation of a stop signal task).

#### Proactive Slowing

Proactive inhibition refers to adjustments made within inhibitory networks during the *expectation* of the potential need to cancel an action (Lavallee et al., 2014). One behavioural manifestation of these changes is proactive slowing. This can be quantified by comparing a participant’s reaction time (RT) to go signals in an experimental condition with no stop trials (i.e., where they are not anticipating any requirement to stop) to their RT in the go trials that occur during a stopping task (van de Laar et al., 2011). Here, RT from the go trials in each of the five SSTs was compared to go trial RT from the go-only block which used the same medium of go signal. The final model used a gamma distribution and log link function (to accommodate for the positive skew in RT data; Lo & Andrews, 2015) and included condition as a single 7-level independent variable (the two go-only blocks with audio or visual go stimuli, and the five SSTs), and age as a covariate. Assessment of proactive slowing in each condition was done with post-hoc tests comparing levels of the condition variable. The random effects structure included participant intercepts and slopes for condition.

#### Behavioural Stopping Delays

Stopping delays are typically assessed by comparing mean RT on successful go trials to the mean RT of the continuing (non-cued) hand in successful stop trials. Because we wanted to compare stopping delays between the various audio/visual cue conditions at the individual trial level we determined a trial-level measure which could be compared across conditions. Each participant’s mean Go trial RT was determined for each condition, and a trial-level stopping delay was calculated by subtracting this from RT of the non-signalled hand in each successful stop trial. The resulting data was analysed using a LMM (Gaussian distribution and identity link function) and included the single factor of condition, with five levels (please refer to Figure 1). The final model used participant intercepts in the random effects structure. Planned Bonferroni-corrected contrasts were conducted to assess effects of modality for both the go and stop stimulus.

### Electromyographic Data Analysis

#### Data Processing

EMG data processing was performed using MATLAB (MathWorks, 2018). Prior to analysis, a fourth-order band-pass Butterworth filter at 20–500 Hz was applied. A single-threshold algorithm was used to determine the precise onset and offset times of EMG bursts (Hodges & Bui, 1996): This involved rectifying and low-pass filtering the data from each trial at 50 Hz before using a sliding window of 500ms to find the trial segment with the lowest root mean squared (RMS) amplitude. This was used as a baseline. EMG bursts were identified where the amplitude of the smoothed EMG was more than 2 SD above this baseline level. If EMG bursts were separated by less than 20ms they were merged to represent a single burst (Weber et al, 2023; Salomoni et al., 2023).

Two types of EMG burst were extracted from the processed data: The *RT-generating burst* was defined as the *last* burst with an onset *after* the go signal but *prior* to the recorded button press. As well as RT-generating bursts, successful stop trials were also assessed for *partial bursts*, i.e., EMG bursts that began decreasing in amplitude before sufficient force was generated to trigger an overt button press. Our algorithm captures such bursts in both the stopping and non-stopping hand. Partial bursts were identified as the earliest burst where peak EMG occurred *after* the presentation of the stop signal but *prior* to the onset of the RT-generating burst from the responding hand. This was done to exclude mirror activity, or other activity unrelated to the task. For partial bursts, peak EMG amplitude was also required to be greater than 10% of the average peak amplitude from that participant’s successful go trials in that condition. For both burst types (RT-generating and partial), we extracted the times of burst onset, peak, and offset (Weber et al. 2023).

EMG envelopes were obtained using full-wave rectification and a low-pass filter at 10 Hz for visual presentation and waveform comparison. For each participant, the amplitude of EMG envelopes from each hand was normalised to the average peak EMG from successful go trials in the go-visual-stop-visual condition. This was done to allow for direct comparisons between participants and conditions.

#### Frequency of Partial Bursts

To determine whether the frequency of partial bursts (i.e., proportion of successful stop trials in which partial bursts were exhibited) differed significantly as a function of the modalities of stop signal, a generalised linear mixed model was run with a binomial distribution and a probit link function. Fixed effects included the single variable of condition, with five levels. The final random effects structure included participant intercepts following convergence issues when a random slope for condition was included.

#### CancelTime

A GLMM was run on data from trials in which a partial burst was detected (see methods section). A single fixed factor was included (condition) with five levels. The final random effects structure included only participant intercepts following issues with model singularity when random slopes were included for condition. As with SSRT, planned Bonferroni-corrected contrasts were used to determine effects of stop modality and go modality, as well as to check for facilitation effects (shortening of CancelTime) in the audiovisual stop signal condition.

#### Action Reprogramming

Response selective stop tasks require the inhibition of the initial bimanual movement followed by rapid action reprogramming to perform a unimanual movement with the non-stopping hand (Gronau et al., 2023). Recently, we have demonstrated that the speed of action reprogramming (defined as the time between the offset of the cancelled bimanual response and subsequent initiation of the RT generating burst in the responding hand on successful selective stop trials) varies between reactive and anticipated movements (Weber et al., 2023b). Given this result in our prior work was not expected, we ran an exploratory analysis here, to check for differences in the speed of action reprogramming in response to auditory and visual stops. A GLMM with a gamma distribution (following identification that the data demonstrated a positive skew similar to that of RT data) and log link function was run with the single fixed factor of condition and a maximal random effects structure to check for differences in the speed of action reprogramming in response to auditory and visual stop stimuli.

## Results

### Behavioural Results

Participants conducted the task well, with high levels of attention indicated by almost 100% accuracy on go trials. Stop success was ∼50%, indicating the SSD staircasing worked well. Summary data is shown in Table 1.

**Table 1:** Key Behavioural Results.

| Condition | Trial Type | RT (ms) | 95%CI (ms) | % Correct | Stopping delays (ms) | 95%CI (ms) | SSRT (ms) | 95%CIs (ms) |
| --- | --- | --- | --- | --- | --- | --- | --- | --- |
| **Visual Go-Only** | Go | 287 | [279 – 294] | 99.8% | - |  | - |  |
| **Auditory Go-Only** | Go | 286 | [278 - 294] | 99.8% | - |  | - |  |
| **Go-Visual-Stop-Visual** | Go<br>Stop | 403<br>511 | [399 – 407]<br>[504 – 518] | 99.8%<br>51.6% | -<br>105 | [90-120] | 206 | [195-218] |
| **Go-Visual-Stop-Audio** | Go<br>Stop | 407<br>533 | [403 – 411]<br>[525 – 541] | 99.6%<br>51.7% | -<br>123 | [108-138] | 200 | [189-212] |
| **Go-Audio-stop-Visual** | Go<br>Stop | 398<br>516 | [394 – 403]<br>[508 – 524] | 99.4%<br>51.1% | -<br>114 | [99-129] | 195 | [184-207] |
| **Go-Audio-Stop-Audio** | Go<br>Stop | 399<br>547 | [395 – 402]<br>[538 – 555] | 99.3%<br>49.6% | -<br>143 | [128-158] | 226 | [214-237] |
| **Go-Visual-Stop-Audiovisual** | Go<br>Stop | 406<br>527 | [402 – 409]<br>[520 – 534] | 99.7%<br>52.8% | -<br>119 | [104-134] | 177 | [165-188] |

*Note: RTs in Go trials represent the average response time of both hands. RTs stop trials represent reaction times of the non-signalled hand (e.g., the right hand RT in a left stop trial). RTs represent descriptive means, stopping delays and SSRTs represent estimated marginal means from the associated model.*

### Proactive Slowing

The proactive slowing analysis revealed a significant main effect of condition χ^2^(6) = 110.15, *p* < 0.001. The covariate of age was not statistically significant χ^2^(1) = 0.547, *p* = 0.46. Bonferroni corrected post-hoc tests revealed that go responses were slower during the SST conditions compared to go only condition using the same medium of go cue, with similar degrees of slowing observed between experimental conditions. Proactive slowing was significant in the Go-Visual-Stop-Visual condition *z* = 8.12, *p* < 0.001, *d* = 1.26, the Go-Visual-Stop-Audio condition, *z* = 9.42, *p* < 0.001, *d* = 1.27, and the Go-Visual-Stop-Audiovisual condition, *z* = 8.68, *p* < 0.001, *d* = 1.29, the Go-Audio-Stop-Audio condition, *z* = 9.18, *p* < 0.001, *d* = 1.20 and the Go-Audio-Stop-Visual condition *z* = 9.43, *p* < 0.001, *d* = 1.10. Post-hoc tests did not reveal any significant differences in go trial RTs between the five experimental conditions (all comparisons *p* = 1.00), suggesting that proactive slowing did not vary across the different experimental conditions.

### SSRT

SSRT analysis revealed a significant main effect of condition *F*(4,112) = 12.88, *p* < 0.001 (Figure 2). Bonferroni corrected contrasts revealed no significant difference between SSRT in auditory stop (Go-Audio-Stop-Audio and Go-Visual-Stop-Audio) and visual stop (Go-Visual-Stop-Visual and Go-Audio-Stop-Visual) conditions, *t*(112) = 2.44, *p* = 0.065, *d* = 0.45. A supplementary Bayesian t-test comparing auditory stops and visual stops demonstrated weak evidence for the null hypothesis BF_01_ = 0.87 (though note strong evidence for this same contrast was observed in the CancelTime analysis). Notably, the SSRTs obtained from conditions with go and stop signals occurring in the *same* modality (Go-Visual-Stop-Visual and Go-Audio-Stop-Audio) were significantly longer than SSRTs in which auditory and visual stimuli were presented in *different* modalities (Go-Visual-Stop-Audio and Go-Audio-Stop-Visual conditions; *t*(112) = 3.43, *p* = 0.002, *d* = 0.75). Furthermore, SSRTs in the Go-Visual-Stop-Audiovisual condition were significantly faster than those in the Go-Visual-Stop-Visual condition, *t*(112) = 4.30, *p* < 0.001, *d* = 1.14 and the Go-Visual-Stop-Audio condition *t*(112) = 3.43, *p* = 0.003, *d* = 0.63.

**Figure 2:**
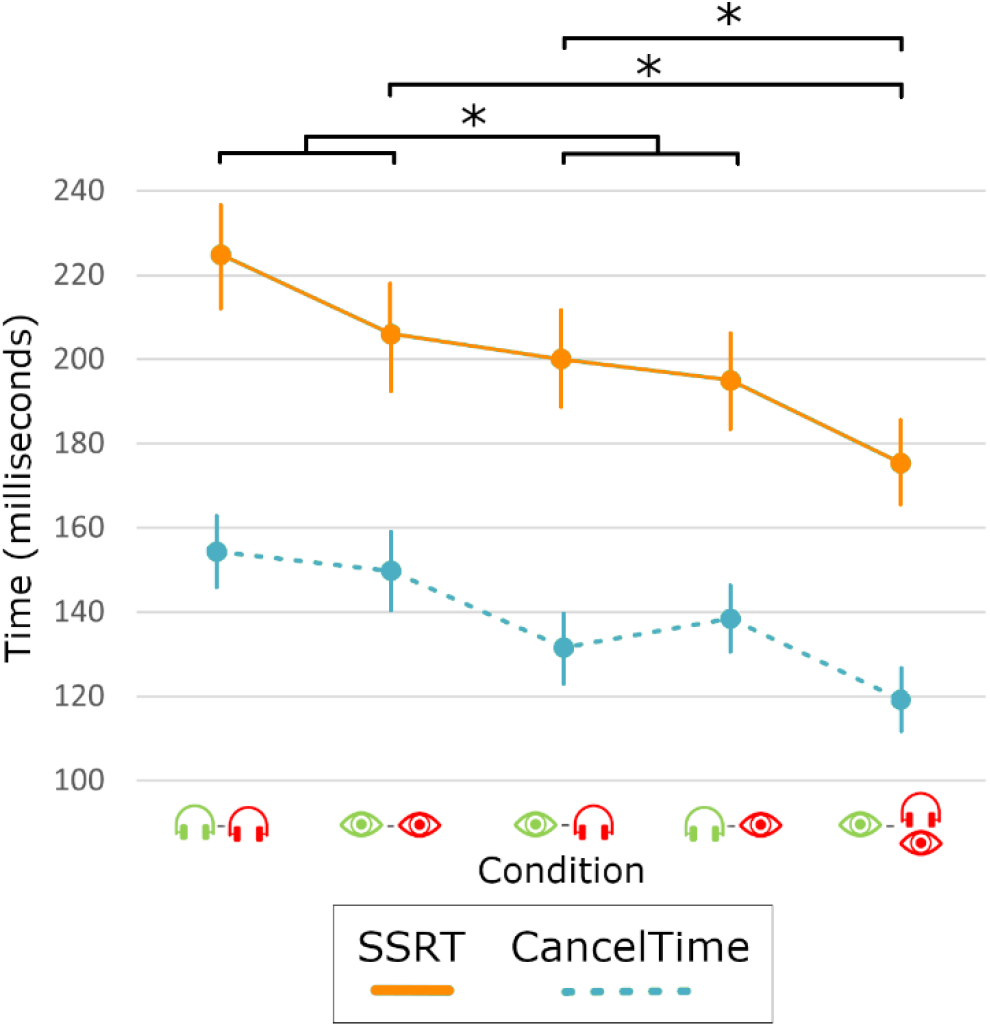
Estimated marginal means of Stop Signal Reaction Time (SSRT) and CancelTime in each of the five conditions. Auditory stimuli are represented with headphones and visual stimuli represented with eyes, with green showing the go stimulus and red showing the stop stimulus. Errors bars represent 95%CIs from the models run on SSRT and CancelTime. Significance flags represent p < 0.01 and apply to both SSRT and CancelTime. The top two significance flags represent multisensory facilitation: The Go-Visual-Audiovisual-Stop condition resulted in faster stopping than the Go-Visual-Stop-Audio and Go-Visual-Stop-Visual condition. The lower (forked) significance flag represents the significant contrasts comparing cross-modal to unimodal tasks.

### Stopping Delays

The results of the GLMM run on stopping delays are depicted in Figure 3a. The model revealed a significant main effect of condition *F*(4,3803) = 23.34, p < 0.001. Planned Bonferroni contrasts revealed that stopping delays in the two conditions with auditory stops were longer than stopping delays in the two conditions with visual stops *t*(3803) = 8.06, *p* < 0.001, *d* = 0.25. Stopping delays in the Go-Visual-Stop-Audiovisual condition were not significantly different those in the Go-Visual-Stop-Audio condition *t*(3803) = 1.07, *p* = 0.851, *d* = 0.05, though they were significantly longer than those in the Go-Visual-Stop-Visual condition *t*(3803) = 3.39, *p* = 0.002, *d* = 0.17.

**Figure 3:**
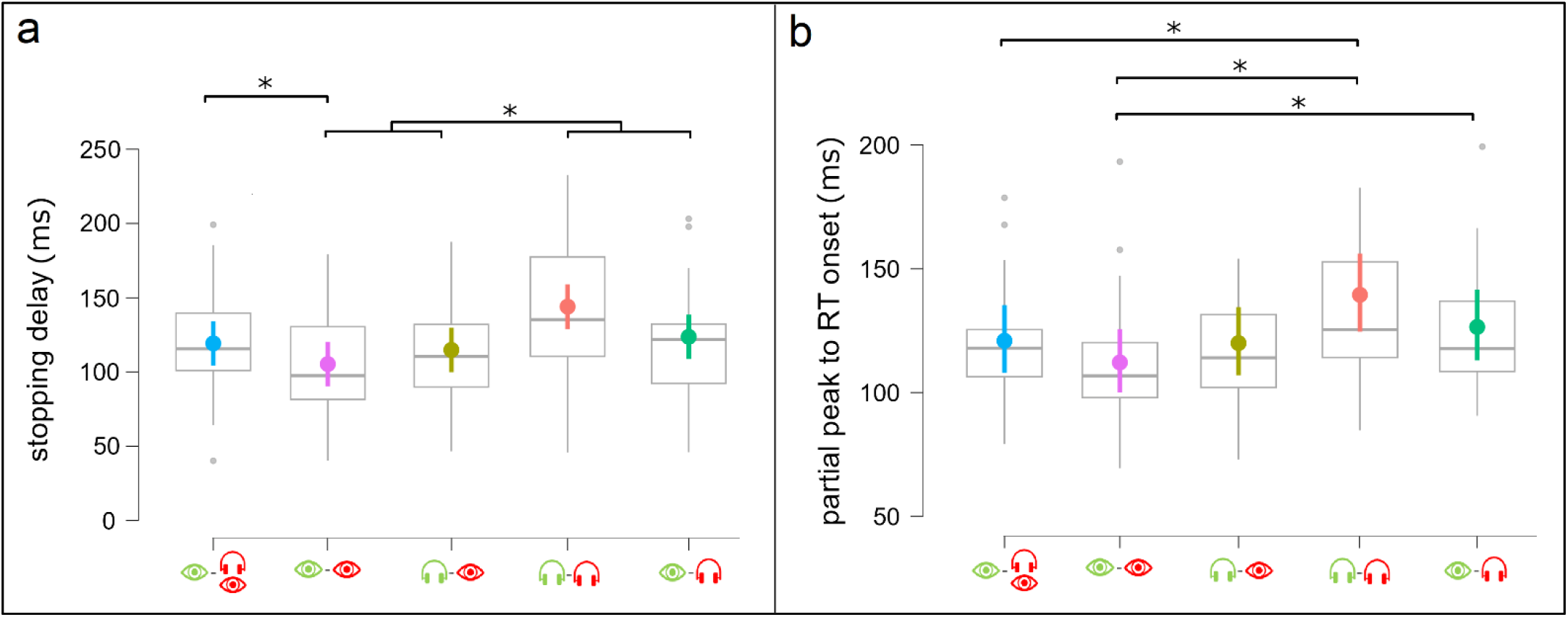
a) central circles represent estimated marginal means from GLMM on stopping delays. b) central circles represent estimated marginal means from GLMM on action reprogramming analysis. * = p < 0.05. Auditory stimuli are represented with headphones and visual stimuli represented with eyes, with green showing the go stimulus and red showing the stop stimulus.

### EMG Results

Table 2 presents the speed of stopping and action reprogramming from the EMG analysis.

**Table 2:** Key Physiological Results.

| Condition | Stop trials with partial bursts | Mean CancelTime (ms) | 95%CI (ms) | Mean Action Reprogramming (ms) | 95%CI (ms) |
| --- | --- | --- | --- | --- | --- |
| **Go-Visual-Stop-Visual** | 37.4% | 149 | [140-159] | 111 | [101-123] |
| **Go-Visual-Stop-Audio** | 50.2% | 132 | [124-140] | 127 | [111-145] |
| **Go-Audio-stop-Visual** | 38.7% | 139 | [130-148] | 120 | [104-139] |
| **Go-Audio-stop-Audio** | 49.0% | 154 | [145-164] | 138 | [121-156] |
| **Go-Visual-Stop-Audiovisual** | 44.7% | 119 | [112-127] | 121 | [109-134] |

### Frequency of Partial Bursts

The GLMM run on presence/absence of partial bursts revealed a main effect of condition χ^2^(4) = 38.52, *p* < 0.001. Planned Bonferroni-corrected contrasts revealed that partial bursts occurred more frequently in the two conditions with auditory stop stimuli compared to the two conditions with visual stop stimuli, *z* = 6.14, *p* < 0.001. When comparing the multisensory stop stimulus condition to the two single sensory stop stimuli conditions with the same visual go stimulus, we observed that Partial bursts were more frequent in the Go-Visual-Stop-Audiovisual condition compared to the Go-Visual-Stop-Visual condition (*z* = 2.82, *p* = 0.015) although no significant differences were noted when this was compared to the Go-Visual-Stop-Audio condition (*z* = 2.08, *p* = 0.11).

### CancelTime

The GLMM conducted on CancelTime revealed a significant main effect of condition χ^2^(4) = 124.07, *p* < 0.001 (Figure 2; Table 2). Planned Bonferroni-corrected contrasts revealed no significant difference between auditory and visual stops when the Go-Audio-Stop-Audio and Go-Visual-Stop-Audio conditions were contrasted with Go-Visual-Stop-Visual and Go-Audio-Stop-Visual conditions(*z* = 0.336, *p* = 1.00, *d* = 0.02). A supplementary Bayesian t-test comparing auditory stops and visual stops confirmed strong evidence for the null hypothesis BF_01_ = 15.78. As with the analysis in SSRT, conditions with go and stop signals occurring in the same modality (Go-Visual-Stop-Visual and Go-Audio-Stop-Audio) demonstrated significantly longer CancelTimes compared to those in which a switch between auditory and visual modalities occurred (Go-Visual-Stop-Audio and Go-Audio-Stop-Visual conditions; *z* = 6.097, *p* < 0.001, *d =* 0.29). CancelTimes in the Go-Visual-Stop-Audiovisual condition were significantly shorter than those in both the Go-Visual-Stop-Visual condition (*z* = 8.13, *p* < 0.001, *d* = 0.70) and the Go-Visual-Stop-Audio condition (*z* = 4.05, *p* < 0.001, *d* = 0.27).

A linear regression, depicted in Figure 4, revealed that SSRTs and mean CancelTimes (i.e., one value per participant, in each condition) demonstrated a moderate positive correlation (*r* = 0.541, *R*^2^ = 0.292, *p* < 0.001).

**Figure 4:**
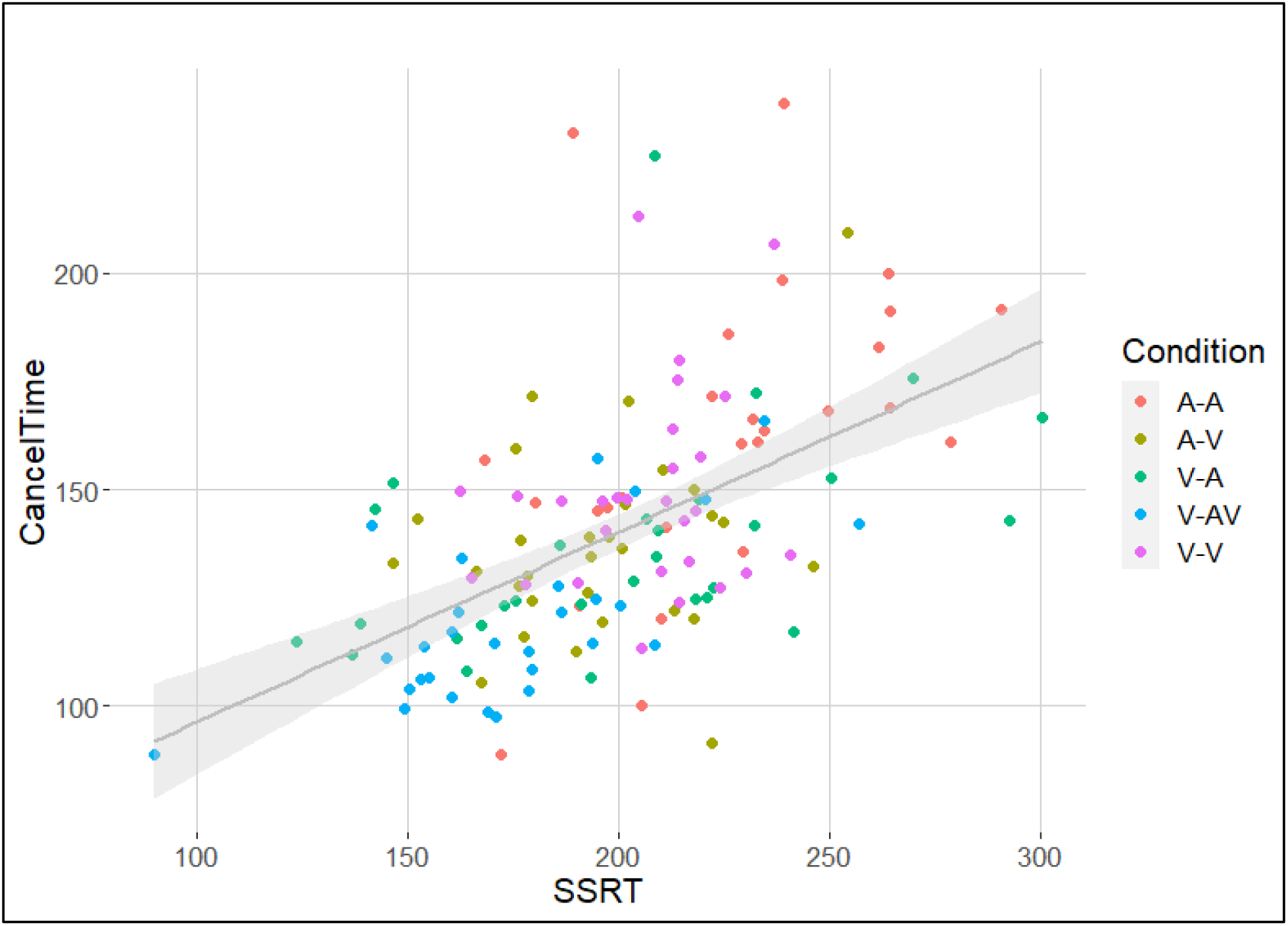
Stop Signal Reaction Time (SSRT) and mean CancelTimes demonstrated a moderate positive correlation. Here, each condition is presented in a different colour, allowing for visualisation of task differences (e.g., SSRTs and CancelTimes for the Go-Visual-Stop-Audiovisual [V-AV; in blue] appear clustered near lower values, while data from the Go-Audio-Stop-Audio condition [A-A; in red] tend towards higher values).

### Action Reprogramming

The results of the GLMM run on action reprogramming are depicted in Figure 3b. The model revealed a significant main effect of condition χ^2^(4) = 26.13, *p* < 0.001 (Table 2 for estimated marginal means). Given that this analysis was exploratory (no specific contrasts were planned), Bonferroni-corrected post-hocs were run on condition (all possible comparisons). Action reprogramming in the Go-Visual-Stop-Visual condition was significantly faster than that in the Go-Audio-Stop-Audio condition (*z* = 5.02, *p* < 0.001, *d* = 0.49), and the Go-Visual-Stop-Audio condition (*z* = 2.98, *p* = 0.03, *d* = 0.35). Action reprogramming was also significantly faster in the Go-Visual-Stop-Audiovisual condition compared to the Go-Audio-Stop-Audio condition (*z* = 3.25, *p* = 0.012, *d* = 0.31). All other comparisons did not reach statistical significance. The full outputs of the post-hoc analysis is presented in Appendix A.

## Discussion

### Stopping latency to stop signals is dependent on go signal modality

The current study used behavioural and physiological measures to compare response-selective stopping performance to auditory, visual, and audiovisual stop signals following different modalities of go signal (auditory or visual). In contrast to the majority of prior research (e.g., Carrillo-de-la-Peña et al., 2019; Morein-Zamir & Meiran, 2003; Ramautar et al., 2006; Van Der Schoot et al., 2005), when conditions using auditory stop stimuli and visual stop stimuli were contrasted, no significant difference was observed in the speed of action cancellation. However, conditions featuring stop and go stimuli in the same modality (go-visual-stop-visual, go-audio-stop-audio) resulted in significantly slower action cancellation than cross-modal conditions (go-visual-stop-audio, go-audio-stop-visual) where a change in stimulus modality occurred between the go stimulus and stop stimulus. As such, the inconsistencies in past research when assessing the effects of stop stimulus modality on stopping speed can be explained by differences in *go stimulus* modality. Specifically, prior research using visual go stimuli observe faster stopping to auditory stop stimuli (Carrillo-de-la-Peña et al., 2019; Morein-Zamir & Meiran, 2003; Ramautar et al., 2006; Van Der Schoot et al., 2005), whereas research using auditory go stimuli do not observe faster stopping in response to auditory stop stimuli (Ikarashi et al., 2022).

One potential explanation for this observation relates to the attentional capacity restrictions which occur when two successive stimuli are presented in the same sensory modality, but not for successive stimuli presented via different modalities (Duncan et al., 1997). That is, separate attentional resources for visual and auditory stimuli (Alais et al., 2006; Wahn & Sinnet, 2019) may have allowed less resource sharing between stop and go processes in the Go-Visual-Stop-Audio and Go-Audio-Stop-Visual conditions, thus resulting in faster cancellations to the stop signal.

The current results challenge the horse-race model of inhibition (upon which traditional measures of SSRT are based; Logan & Cowan, 1984), which assumes that stop and go processes operate independently. It follows that an independent stop processes, triggered uniquely by the stop signal, should not be influenced by the modality of the go signal, as was observed in the current research. Notably, past research assessing the independence of stop and go processes has used cross-modal tasks. For example, research by Yamaguchi and colleagues (2011) sought to determine whether the “psychological refractory period effect” (which describes how, when enacting responses to two stimuli occurring in rapid succession, the response to the second stimulus is delayed; Levy et al., 2006; Lien et al., 2006; Pashler, 1994) applied in stop signal paradigms. They found that this phenomenon did not apply when the second “response” was inhibitory in nature (that is, there was no effect on SSRT), and argued that this was further evidence of independence between go and stop processes. Similar results have been observed in earlier research (Logan and Burkell, 1986). Notably though, both of these experiments employed visual go signals and auditory stops. It is plausible then, that in a unimodal task, the psychological refractory period does influence the stop process, contributing to the current result. A within subject comparison of the delays associated with two consecutive go tasks (i.e., a dual task paradigm; Dux et al., 2006; Wahn & Sinnet, 2019) and the delays in the stop process, in unimodal and cross-modal tasks, could shed further light on this issue.^2^ Further research using neuroimaging or electroencephalographic techniques in combination with different modalities of go and stop stimuli would provide further insights into this theoretical perspective.

### Multisensory stimulation improves stopping speed

We observed that a stop signal featuring simultaneous visual and auditory components resulted in a faster inhibitory response than unimodal stop signals (visual or auditory; Figure 2). This novel finding suggests that a multisensory stop stimulus (Hershenson, 1962; Molholm et al., 2002) can facilitate inhibitory responses, and contrasts with recent research observing *slowed* inhibitory responses to multimodal stop signals (Strelnikov et al., 2021). This difference in results likely relates to aspects of task design. Notably, the research by Strelnikov et al. sought to increase ecological validity, using human faces as visual stimuli and human voices as auditory stimuli. Given the unique effects of faces on human attentional systems (Palermo & Rhodes, 2007), these results may be specific to the stimuli used in that study. Alternatively, it may be that multisensory facilitation enhances inhibition when simple stimuli are used (e.g., simple coloured circles and sine waves, as in the current experiment) but not when more complicated stimuli are used (e.g., complex images, recognisable sounds). Further research using both simple, complex, and face-based stimuli could provide clarification on this point. We note also that research by Stock et al. (2017) did not observe a significant difference in SSRTs when comparing visual, audiovisual, and tactile-visual cues in a stop-change paradigm. The reason for this finding is less clear, though likely also relates to task complexity, as their experiment involved more complex responses and response mappings (i.e., the relation between the stimulus and the associated response).

### Auditory stop signals result in more partial bursts and longer selective stopping delays

The percentage of successful stop trials in which partial bursts were elicited (i.e., muscle activation was initiated but cancelled prior to eliciting an overt button press) was significantly greater in response to auditory stop signals than visual stop signals (refer to Table 2). The reason for this is unclear, though as a practical consideration, if researchers are primarily seeking to calculate inhibition latency using EMG (Raud et al., 2022), auditory stop stimuli could be considered to improve validity of the result due to the greater expected proportion of partial bursts., thus reducing the censoring effects of only analysing a subset of stop trials (see Salomoni et al. 2023).

Interestingly, although stopping speed depended on the combination of go and stop signal modality, stopping delays (i.e., the delay in the ongoing action following successful stopping) were longer following auditory stops, irrespective of go signal modality. This result is consistent with previous research using a stop-change paradigm, which observed slower RTs following a stop-change stimulus in the auditory modality compared to a visual stop-change stimulus (Stock et al., 2017). Stock et al. also noted larger event-related potentials related to conflict and cognitive effort (i.e., central P3 amplitudes) following auditory stop-change stimuli than visual stop-change stimuli. These observations likely relate to differences in processes involved in translating the location of auditory and visual stimuli into motor responses (Bushara et al., 1999). In the current task, difficulties related to response mapping between the monophonic auditory stimuli and the requisite buttons may explain the larger stopping delays in auditory stop conditions. That is, cognitive processes involved in identifying the side of the tone may have taken longer than those involved in identifying the side of the colour change. Notably, sound identification and sound localisation rely on distinct auditory streams (Alain et al., 2001). As such, comparing the method used in the current study to a future selective stopping task which trained participants to use pitch to differentiate response mappings (e.g., high frequency = le; low frequency = right) may find a different result. In sum, this finding provides evidence that complex stopping tasks are subject to differences in sensory modality (i.e., more efficient selective stopping is possible to environmental cues in the visual modality than the auditory modality), though further research is required to assess the generalisability of this finding beyond the current experimental paradigm.

### Conclusions

The current research provides novel evidence that stopping speed to auditory and visual stop signals is dependent on the modality of the preceding go signal. Furthermore, we report the novel finding of multisensory facilitation of the action cancellation process, suggesting that even action stopping – thought to be subserved by rapid cortical-subcortical pathways – is modifiable. This research emphasises the importance of sensory modality in inhibitory research, highlighting how domain-specific processes interact in a complex manner with task design, influencing observations in the wide variety of clinical and non-clinical research domains which use the stop signal task. More broadly, these findings bear implications for human-factors engineering, especially in contexts which require the rapid coordination of limbs (i.e., implementation of beeps vs visual displays warnings in vehicles).

## Data availability statement

The data that support the findings of this study are available from the corresponding author upon reasonable request.

## Author contributions

**Simon Weber:** conceptualisation, methodology, software, writing – original draft, formal analysis, investigation. **Sauro Salomoni:** conceptualisation, methodology, software, formal analysis, writing – review & editing. **Rebecca StGeorge:** conceptualisation, writing – review & editing, supervision. **Mark Hinder:** conceptualisation, resources, writing – review & editing, supervision, project administration, funding acquisition.

## Appendix 1 Action reprogramming post-hoc tests

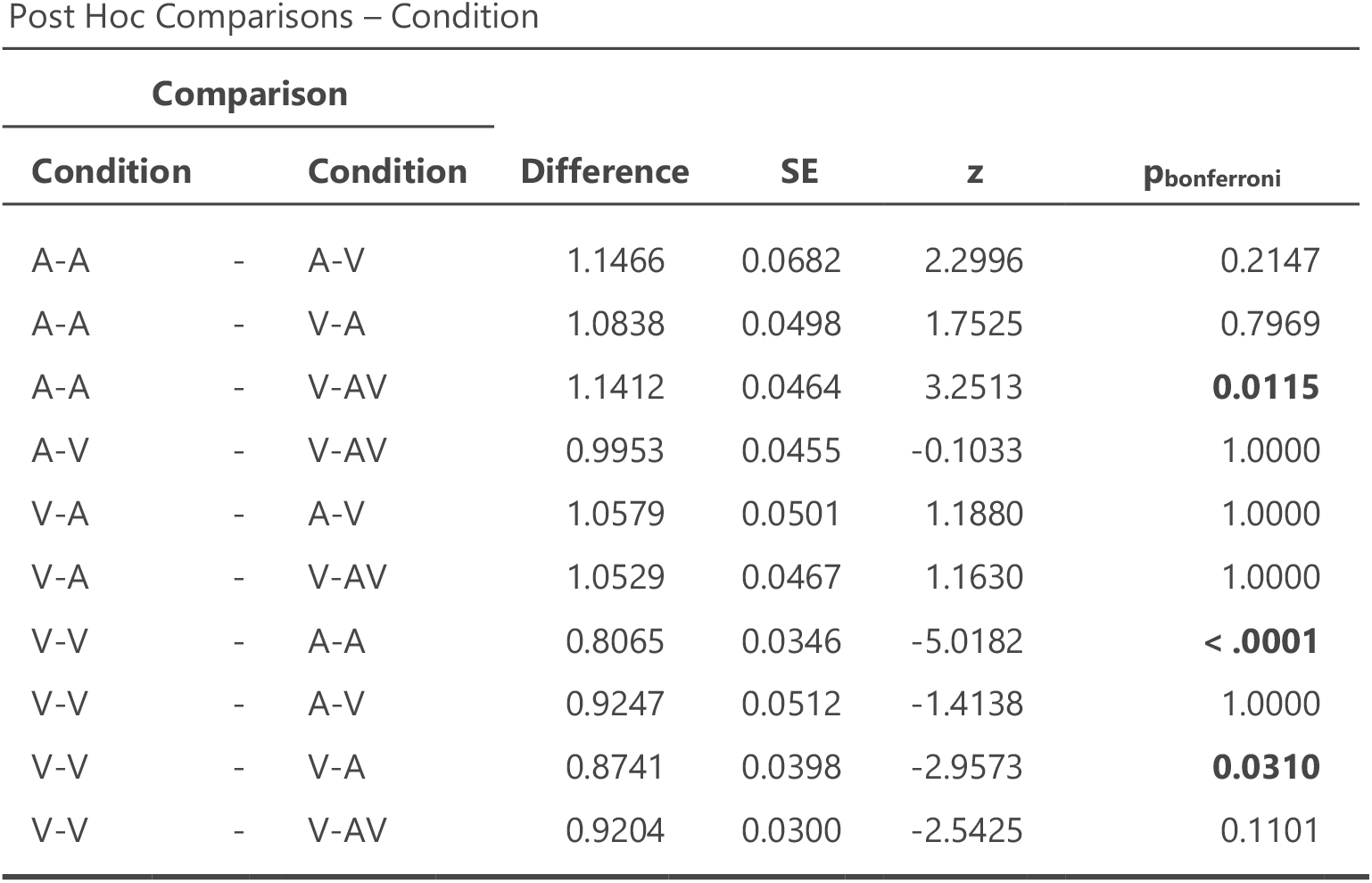

## Footnotes

1 In the Go-Audio-Stop-Audio condition if the stop signal started before the go-tone had finished playing (i.e., at SSDs < 200ms), the tone of the go cue (220Hz, both headphones) was stopped at the same time that the stop cue (440Hz, one headphone) began, to prevent the tones from playing simultaneously.

2 Though we note that the psychological refractory period exists in both unimodal and cross-modal dual-tasks (Morrison et al., 2015; Spagna et al., 2020). Thus, an argument could be made that it’s absence in a cross-modal stopping task renders its absence in a unimodal task unlikely. To the authors knowledge, no research has directly investigated this.

